# Co-targeting menin and LSD1 dismantles oncogenic programs and restores differentiation in MLL-rearranged AML

**DOI:** 10.1101/2025.10.13.681683

**Authors:** Mina M. Tayari, Helena Gomes Dos Santos, Felipe Beckedorff, Tulasigeri Totiger, Adnan Mookhtiar, Anna Lesley Kingham, Guilherme Miura Lavezzo, Gloria Mas Martin, Hsuan-Ting Huang, Efe Karaca, Daniel Bilbao, Justin Taylor, Ramin Shiekhattar, Justin M. Watts

**Author notes:** correspondence to: Justin Taylor, MD, Ramin Shiekhattar, PhD, Justin Watts, MD. Equal contribution (Justin Taylor, Ramin Shiekhattar, Justin M. Watts).

## Abstract

Acute myeloid leukemia (AML) harboring *MLL* (*MLL1, KMT2A*) rearrangement (*MLL-r*) remains a lethal subtype with limited durable responses to single-agent menin inhibition. To define rational combination strategies, we performed a high-throughput screen of >900 epigenetic modulators in combination with menin inhibition in *MLL-r* AML models. This uncovered consistent synergy between menin and lysine-specific demethylase 1 (LSD1) inhibition, including with the clinical agent iadademstat. Mechanistically, LSD1 was found to interact with LEDGF/p75 (PSIP1), a chromatin-anchoring cofactor of the menin-MLL complex at H3K36me3 marked euchromatin. Chromatin profiling revealed extensive co-occupancy of LSD1 and menin-MLL components at leukemogenic loci in MLL-r AML cells. Dual inhibition of menin and LSD1 dismantled this chromatin complex, evicted H3K36me3 from LEDGF-bound sites, and reprogrammed transcription toward myeloid differentiation. Combined menin and LSD1 blockade repressed canonical MLL targets, including *HOXA9, MYC, FLT3, PBX3*, and *CDK6*, while restoring H3K36me3 and H3K4me3 and activating differentiation-associated genes. In vivo, the combination produced potent antileukemic effects in both MOLM-13 and *MLL-r* patient-derived xenografts, markedly reducing leukemic burden and extending survival without overt toxicity. These findings identify LSD1 as a critical cofactor of the menin-MLL-LEDGF axis and establish concurrent menin and LSD1 inhibition as a mechanistically informed combinatorial therapeutic approach in *MLL-r* AML.

## INTRODUCTION

Chromosomal translocations involving mixed lineage leukemia rearrangement (*MLL-r*) generate fusion proteins that aberrantly regulate gene expression and drive leukemogenesis [1-3]. MLL-r define a high-risk subtype of acute myeloid leukemia (AML), associated with poor prognosis and frequent relapse [2, 4, 5]. *MLL-r* occur in approximately 10% of leukemias [6-8], accounting for 15–20% of pediatric AML and up to 50% of infant cases, and are associated with poor prognosis compared to non *MLL-r* disease [2, 5, 9]. Standard therapies often fail to achieve durable remission, underscoring the urgent need for novel treatments. Menin inhibitors have emerged as promising therapeutics in relapsed/refractory (R/R) *MLL-r* AML with revumenib approval in November 2024, marking it the first-in-class menin inhibitor approved to date. However, the response duration in the AUGMENT-101 trial averaged just over 6 months [10]. Here, we hypothesized that combining menin inhibitors with other epigenetic-modifying drugs would synergistically disrupt leukemogenic transcriptional programs and enforce stable differentiation, thereby yielding more durable therapeutic responses in *MLL-r* AML.

Menin interacts directly with lens epithelium-derived growth factor LEDGF/p75 (PSIP1) [11-13]. Disruption of LEDGF in murine models impairs MLL fusion-mediated leukemogenesis and reduces Hox gene expression, underscoring its essential role in stabilizing the oncogenic transcriptional machinery [13-15]. Together, the menin-MLL-LEDGF ternary complex enforces transcriptional programs that enhance proliferation and block hematopoietic differentiation [1, 11, 13]. The N-terminal region of MLL and MLL fusion proteins interacts with menin and LEDGF, which together anchor the complex to chromatin and maintain its association with target loci [11, 13, 14]. LEDGF harbors a methyl-lysine-binding PWWP (Pro-Trp-Trp-Pro) domain that recognizes H3K36me2/3 at actively transcribed regions and functions as a chromatin-tethering coactivator, anchoring interacting proteins to H3K36me2/3-enriched loci [16-18]. Genetic and biochemical studies demonstrate that both menin and LEDGF are essential for MLL fusion-driven leukemogenesis, whereas LEDGF appears largely dispensable for normal hematopoiesis [19]. Notably, menin is also required for leukemogenesis driven by nucleophosmin 1 (*NPM1)* mutations, broadening its therapeutic relevance [20]. Pharmacologic inhibition of the MLL-menin interaction has emerged as a promising therapeutic strategy [7, 10]. Preclinical studies demonstrate that menin inhibitors attenuate MLL-fusion-driven transcriptional activation of genes essential for hematopoietic cell proliferation, thereby restoring differentiation and reducing leukemic burden [7, 21]. In early-phase clinical trials, the oral menin inhibitor revumenib (SNDX-5613) produced molecular responses and reduced leukemic blasts in patients with *MLL-r* and *NPM1*-mutated AML [22-24]. Menin inhibition also synergizes with other targeted therapies, providing a strong rationale for combination approaches, though currently only FDA approved as a monotherapy.

In this study, we screened for epigenetic-modifying compounds that enhance the potency of menin inhibition. Lysine-specific demethylase 1 (LSD1) emerged from this screen as a particularly compelling combinatorial partner. LSD1 (KDM1A) is a flavin-dependent histone demethylase that functions as an epigenetic eraser, removing activating H3K4me1/2 marks through the CoREST complex to maintain transcriptional repression [25, 26]. Beyond its enzymatic role, LSD1 shapes chromatin accessibility and sustains leukemic stemness by blocking myeloid differentiation. [27-29]. Prior clinical studies with the nonselective LSD1 inhibitor tranylcypromine (TCP) demonstrated activity in AML when combined with all-trans retinoic acid (ATRA) [30, 31]. However, its clinical use was limited by dose-dependent toxicity and off-target inhibition of monoamine oxidase A (MAO-A). Next-generation, selective LSD1 inhibitors such as iadademstat (ORY-1001) overcome these limitations and are under active clinical evaluation in AML [32].

To study the mechanisms of the synergy of dual menin and LSD1 inhibition, we used MLL-AF9-rearranged MOLM-13 and MLL-AF4-rearranged MV4-11 AML cell lines, together with patient-derived xenograft models. We demonstrate that the combination disrupts the menin-MLL-LEDGF transcriptional activity, represses canonical leukemogenic targets, and restores expression of differentiation-associated genes. These findings highlight a mechanistically informed therapeutic strategy with clear translational relevance for MLL-r AML and provide strong rationale for clinical testing of menin-LSD1 inhibitor combinations in MLL-r acute leukemia.

## RESULTS

### High-throughput epigenetic screening identifies LSD1 inhibition as a synergistic vulnerability with menin blockade in MLL-r AML

We performed a high-throughput screen (HTS) of 932 epigenetic modulators to identify compounds that synergize with menin inhibition in MLL-r AML cell lines. The screen validated previously reported synergy between the menin inhibitor DSP-5336 and targets shown to synergize with other menin inhibitors, such as BRD4, DOT1L and KAT6A [33-35] **(Fig. 1A**). Notably, all seven LSD1 inhibitors tested, including the clinical compound ORY-1001, enhanced cell death when combined with menin inhibition, resulting in decreased viability in MV4-11 cells (**Fig. 1B**). We next evaluated the combination of the menin inhibitor SNDX-5613 with the LSD1 inhibitor ORY-1001 in MV4-11 (MLL-AF4-rearranged) and MOLM-13 (MLL-AF9-rearranged) cells. In MV4-11 cells, which were highly sensitive to menin inhibition alone (IC_50_ = 40 nM for SNDX-5613), ORY-1001 showed an IC_50_ of 1.67 μM. The combination reduced the IC_50_ to 8.39 nM, indicating synergetic effects (**Fig. 1C**). In contrast, MOLM-13 cells showed only modest viability changes with SNDX-5613 alone (IC_50_ = 1.62 μM) and, consistent with prior observations that MOLM-13 is resistant to LSD1 inhibition [29], exhibited limited response to ORY-1001. However, the combination dramatically decreased the IC_50_ to 14.9 nM (**Fig. 1D**). Fluorescence-activated cell sorting (FACS) analysis revealed an increased percentage of CD11b^+^ cells with the combination treatment, indicative of enhanced differentiation relative to single agents (**Fig. 1E**).

**Figure 1.**
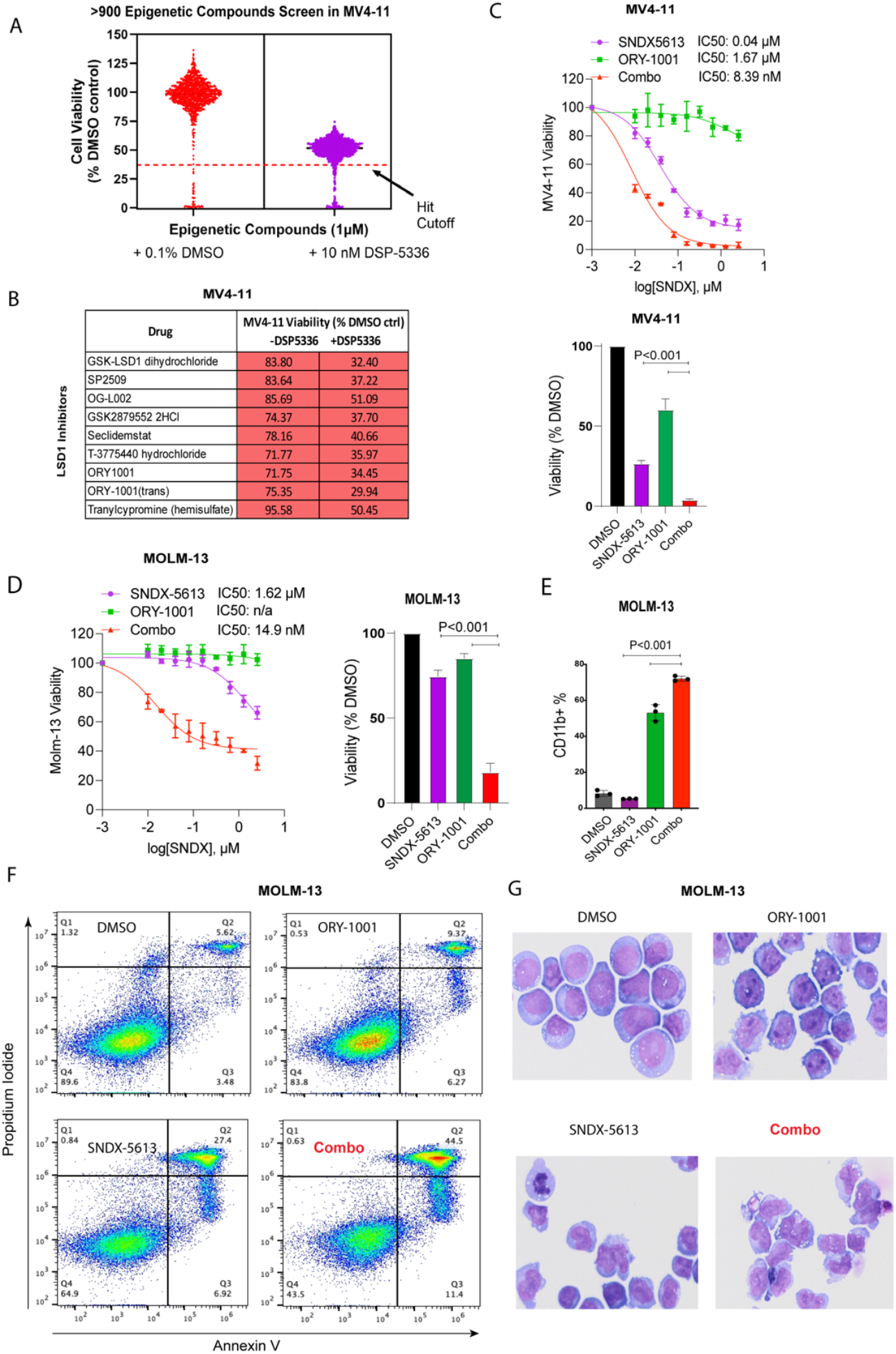
Combined LSD1 and menin inhibition synergistically suppresses AML cell proliferation. (**A**) High-throughput screen (HTS) of 932 epigenetic modulators in MV4-11 cells with the menin inhibitor DSP-5336. (**B**) Cell viability of MV4-11 cells treated with DSP-5336 in combination with seven LSD1 inhibitors, including ORY-1001. (**C**) Dose-response curves and IC_50_ values for SNDX-5613, ORY-1001, and their combination in MV4-11 (MLL-AF4) and (**D**) MOLM-13 (MLL-AF9). n/a, not applicable; IC_50_ undetermined. (**E**) Percentage of CD11b^+^ cells measured by fluorescence-activated cell sorting (FACS) after single and combination treatments. (**F**) Flow cytometry analysis of Annexin V/7-AAD staining in MOLM-13 cells (**G**) May-Grünwald-Giemsa-stained cytospin preparations illustrating morphological changes in MOLM-13 cells treated with DMSO, SNDX-5613, ORY-1001, or their combination.

To determine whether the combination induces apoptosis in MOLM-13 cells, we performed flow cytometry with Annexin V/7-AAD staining. Treatment with ORY-1001 or SNDX-5613 alone increased late apoptotic cells to 9.37% and 27.4%, respectively, compared to 5.62% in controls. However, the combination further elevated late apoptosis to 44.5% (**Fig. 1F**). The observed phenotype was consistent with cell differentiation, as cytospin and morphological analyses revealed enhanced differentiation-associated changes with the combination treatment (**Fig. 1G**). Collectively, these results indicate that dual inhibition of menin and LSD1 promotes differentiation and apoptosis, producing more potent antileukemic activity than single-agent treatment.

### RNA-seq reveals synergistic transcriptomic reprogramming by combined menin and LSD1 inhibition

We performed RNA sequencing (RNA-seq) to profile transcriptomic changes in MOLM-13 and MV4-11 cells after 24 hours of treatment with DMSO (control), ORY-1001 (0.2 µM), SNDX-5613 (0.2 µM), or the combination. In MOLM-13 cells, ORY-1001 treatment resulted in 36 genes downregulated and 280 genes upregulated, whereas SNDX-5613 altered 180 genes down and 307 genes up. Combination treatment produced the most profound effect, with 966 genes downregulated and 569 upregulated (**Fig. 2A**). MV4-11 cells exhibited greater transcriptomic sensitivity compared to MOLM-13, consistent with their enhanced sensitivity in proliferation assays. ORY-1001 alone caused 51 downregulated and 386 upregulated genes, largely consistent with LSD1 inhibition leading to derepression of target genes. SNDX-5613 treatment had a more dramatic effect, with 986 genes downregulated and 1308 upregulated. Combination treatment further amplified these effects, yielding 1761 downregulated and 1586 upregulated genes (**Fig. 2B**). While SNDX-5613 already induced broad transcriptomic changes as a single agent, the combination produced an even stronger effect. In contrast, MOLM-13 cells displayed more balanced responses to each single agent, with the combination producing a distinct transcriptional signature. Notably, in MOLM-13 cells, 942 genes were uniquely regulated by the combination treatment **(Fig. 2C)**, while in MV4-11 cells, 1,317 genes were uniquely regulated **(Fig. 2D)**, underscoring the shared yet enhanced transcriptional reprogramming induced by combined LSD1 and menin inhibition.

**Figure 2.**
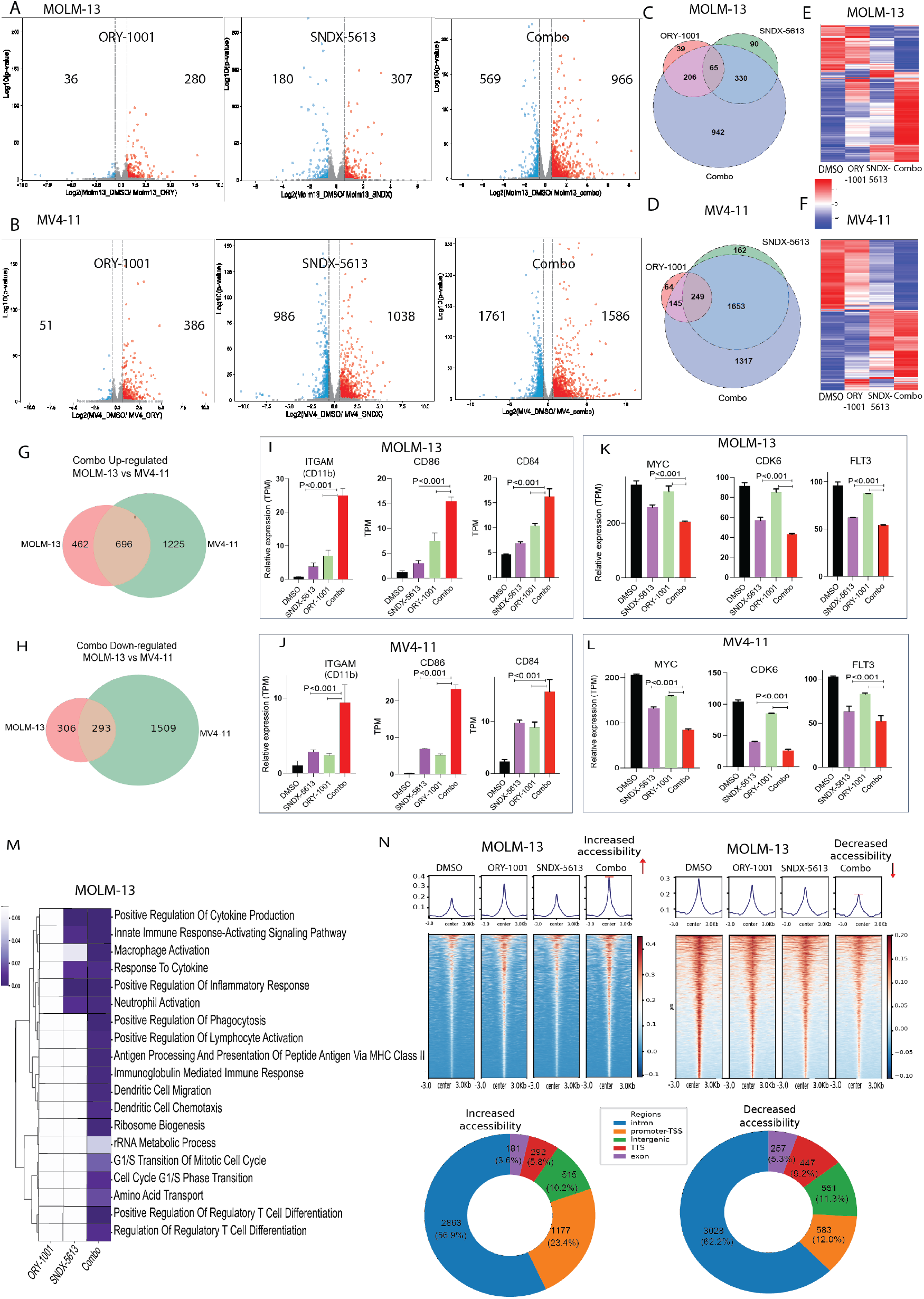
Transcriptomic reprogramming induced by combined menin and LSD1 inhibition. **(A)** Volcano plots of differentially expressed genes (DEGs) in MOLM-13 and **(B)** MV4-11 after 24 h treatment with DMSO, ORY-1001 (0.2 µM), SNDX-5613 (0.2 µM), or the combination. (**C**) Venn diagram showing unique and shared DEGs across treatment groups in MOLM-13 and **(D)** MV4-11. (**E**) Heatmap illustrating synergistic up- and downregulation of genes, with combination treatment amplifying transcriptional changes observed with single agents in MOLM-13 and **(F)** MV4-11. (**G**) Overlap of up- and downregulated genes in MOLM-13 and (**H)** MV4-11 following combination treatment. (**I-J**) Induction of differentiation markers (*CD11b, CD86*) in MOLM-13 and MV4-11 cells. (**K-L**) Downregulation of MLL target oncogenes *MYC, CDK6* and *FLT3*, in both cell lines. (**M**) Pathway enrichment analysis showing strong induction of innate and adaptive immune activation, cell cycle regulation, and metabolic processes with the combination. (**N**) ATAC-seq analysis demonstrating synergistic increased or decreased chromatin accessibility with combination treatment compared to either single agent.

Heatmap analysis revealed that LSD1 inhibition induced modest transcriptional changes, whereas menin inhibition produced a more pronounced effect, with the combination treatment yielding the strongest overall response **(Fig. 2E-F)**. In MOLM-13 cells, genes that were highly expressed in the DMSO control showed a stepwise decrease in expression with LSD1 inhibition, a further decrease with menin inhibition, and the most substantial downregulation with the combination. Conversely, genes with low basal expression in DMSO were modestly upregulated by either LSD1 or menin inhibition alone, with synergistic upregulation observed upon combined treatment **(Fig. 2E)**. A similar pattern was observed in MV4-11 cells, where LSD1 inhibition produced minor effects, menin inhibition drove the predominant transcriptional response, and the combination treatment elicited the strongest modulation across both up- and downregulated gene sets **(Fig. 2F)**. Overall, heatmap analysis showed that genes modestly affected by either single agent were further altered in the same direction with the combination, indicating additive or synergistic transcriptional effects across most genes.

Although MOLM-13 cells exhibited overall fewer transcriptional changes in response to the combination, more than half of the genes that were up- or downregulated still overlapped with those altered in MV4-11 cells, indicating shared combination-specific transcriptional programs (**Fig. 2G-H**). Notably, genes associated with myeloid differentiation, including *CD11b, CD86*, and *CD84*, showed modest upregulation with single-agent treatment and were dramatically increased with the combination in both cell lines **(Fig. 2I-J)**. In contrast, key oncogenic drivers regulated by MLL, including *MYC, FLT3* and *CDK6* (**Fig. 2 K-L**), *HOXA9, Meis1* (**Fig. S2)**, showed modest downregulation with single-agent treatment and were markedly suppressed by the combination in both MOLM-13 and MV4-11 cells. Pathway enrichment analysis further revealed that the combination, compared to either single agent, elicited strong immune activation signatures (innate and adaptive), alongside regulation of cell cycle and metabolic processes, indicative of broad transcriptional reprogramming (**Fig. 2M**). ATAC-seq analysis demonstrating synergistic alterations in chromatin accessibility, with both increases and decreases observed upon individual drug treatment, and the combination producing the most pronounced gains or losses across both cell lines (**Fig. 2N** and **Fig. S1**).

### Immunoprecipitation mass spectrometry of endogenous LSD1 identifies previously unrecognized interaction with LEDGF

To determine whether LSD1 could be interacting directly with menin to explain the synergy of the combined inhibition of these targets, we performed immunoprecipitation of LSD1 in Molm-13 cells followed by mass spectrometry. This analysis confirmed canonical LSD1-associated partners, including HDAC1/2 and RCOR1/3 [26, 36-38], as well as previously reported interactors GFI1 [39], GSE1 [40, 41] and CARM1 [42]. Notably, while menin was not among the proteins co-immunoprecipitated with LSD1, the analysis revealed a previously unrecognized association with LEDGF (**Fig. 3A**).

**Figure 3.**
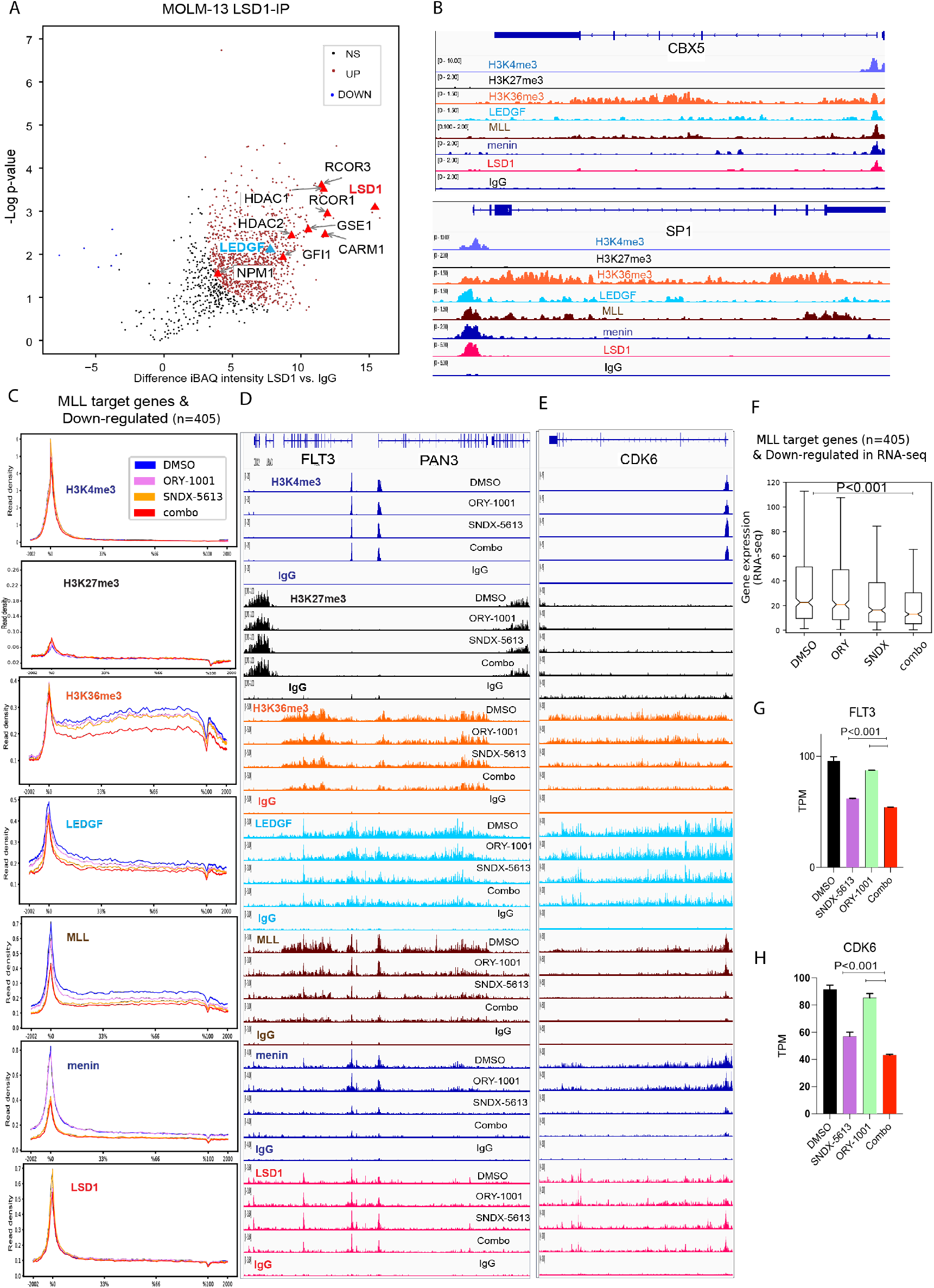
Dual inhibition of LSD1 and menin disrupts MLL complex occupancy and promotes differentiation. (**A**) Mass spectrometry of endogenous LSD1-IP reveals a novel interaction with LEDGF. **(B)**CUT&RUN shows co-occupancy of LSD1, MLL, Menin, and LEDGF at representative loci (*CBX5, SP1*). (**C**) Global metaplot of associated changes in H3K4me3, H3K27me3, H3K36me3, LEDGF, MLL, Menin, and LSD1 in MLL target genes (n = 405), that are downregulated by RNA-seq. (**D-E**) Genome tracks at MLL targets (*FLT3* and *CDK6*). (**F)** Box plot of downregulated MLL target genes following treatment with DMSO, SNDX-5613, or the combination treatment. **(H)** TPM (transcripts per million) RNA expression of *FLT3* and *CDK6* in cells treated with DMSO, SNDX-5613, or the combination treatment. Statistical analysis was performed using one-way ANOVA followed by Tukey’s post hoc test (single treatments vs. combination).

This interaction suggests a direct mechanistic link between LSD1 and the menin-MLL-LEDGF axis, a transcriptional coactivator that harbors an H3K36me3-recognizing PWWP domain and is known to bridge menin and the N-terminus of MLL proteins. To our knowledge, this represents the first demonstration of an LSD1-LEDGF association.

To further probe the chromatin context of this interaction, we performed CUT&RUN profiling, which revealed substantial co-occupancy of LSD1 with MLL complex components across the genome. Representative CUT&RUN tracks at the *CBX5* and *SP1* loci show coordinated enrichment of the MLL complex, menin, and LEDGF, and reveal LSD1 co-occupancy at the promoters, while the gene bodies are enriched for the H3K36me3 activation mark (**Fig. 3B**). As expected, LEDGF occupancy correlated with H3K36me3, while repressive H3K27me3 was depleted at these loci, indicating localization to transcriptionally broad regions of euchromatin. **Fig. 3C** shows global analysis of MLL target genes that are downregulated by RNA-seq. Reduced H3K36me3 is accompanied by loss of the MLL and LEDGF at the same region in this subset of genes bound by MLL and downregulated with combination (**Fig. 3C)**. At *FLT3-PAN3*, H3K4me3 decreased modestly with the combination, and H3K27me3 flanked H3K36me3-marked active gene bodies (**Fig. 3D**). In agreement with prior work by Leroy et al. [18], we observe a pronounced opposition between H3K27me3 and H3K36me3 across the genome: H3K27me3 is depleted over H3K36me3-enriched gene bodies yet accumulates in flanking regions, demarcating the boundaries of active transcription (Fig. 3D).

As shown by the metaplot (**Fig. 3C**) and at representative MLL targets, *FLT3-PAN3, CDK6* (**Fig. 3D-E**), and the *HOX* cluster, *MEIS1*, and *PBX3* (**Fig. S2**) treatment with the LSD1 inhibitor ORY-1001 alone unexpectedly reduced MLL and LEDGF occupancy and diminished H3K36me3, to a degree comparable to menin inhibition. Menin blockade with SNDX-5613 further decreased MLL and LEDGF chromatin binding, and the combination exacerbated these effects, culminating in near-complete eviction of H3K36me3 from LEDGF-bound loci.

Menin occupancy was selectively reduced by menin inhibition, with little to no change between DMSO and LSD1 inhibition, and no appreciable further decrease upon combination treatment (**Fig. 3C-D**). Interestingly, LSD1 chromatin association appeared slightly increased in SNDX-5613-treated cells, suggesting compensatory stabilization of LSD1 at these loci in the absence of menin-MLL. Collectively, these data support a model in which LSD1 and LEDGF cooperate at transcriptionally active chromatin, but disruption of menin-MLL severely compromises H3K36me3 maintenance, revealing a previously unappreciated role for LSD1 in the structural and functional integrity of this complex.

The CUT&RUN-defined set of 405 MLL-bound targets exhibited reduced expression with ORY-1001 or SNDX-5613, with a more pronounced decrease under the combination **(Fig. F)**. Representative RNA expression of *CDK6* and *FLT3* further demonstrated decreased levels upon single treatment, which were further reduced with the combination **(Fig. G-H)**.

### Synergistic effects of inhibitors of menin and LSD1 induce genes critical for differentiation

The effects of dual inhibition extended beyond repression of canonical MLL targets oncogenic loci. We and others have shown that LSD1 function as a barrier to myeloid differentiation and enhancers of lineage-specifying genes [30, 31, 43, 44]. Consistent with this role, we observed that the combination of ORY-1001 and SNDX-5613 not only disrupted MLL complex occupancy at leukemogenic genes but also promoted epigenetic changes at differentiation-associated loci. Among the CUT&RUN-defined MLL-bound genes, we analyzed the subset upregulated by treatment in RNA-seq relative to DMSO (n = 413). This subset of genes exhibited increased H3K36me3 enrichment across the gene body upon treatment. Interestingly, treatment with ORY-1001 and SNDX-5613 each caused an increase as single agents; however, the most prominent increase was observed when the treatments were combined **(Fig. 4A)**, and this was further quantified in the box plot showing significant changes after combination treatment **(Fig. 4B)**.

**Figure 4.**
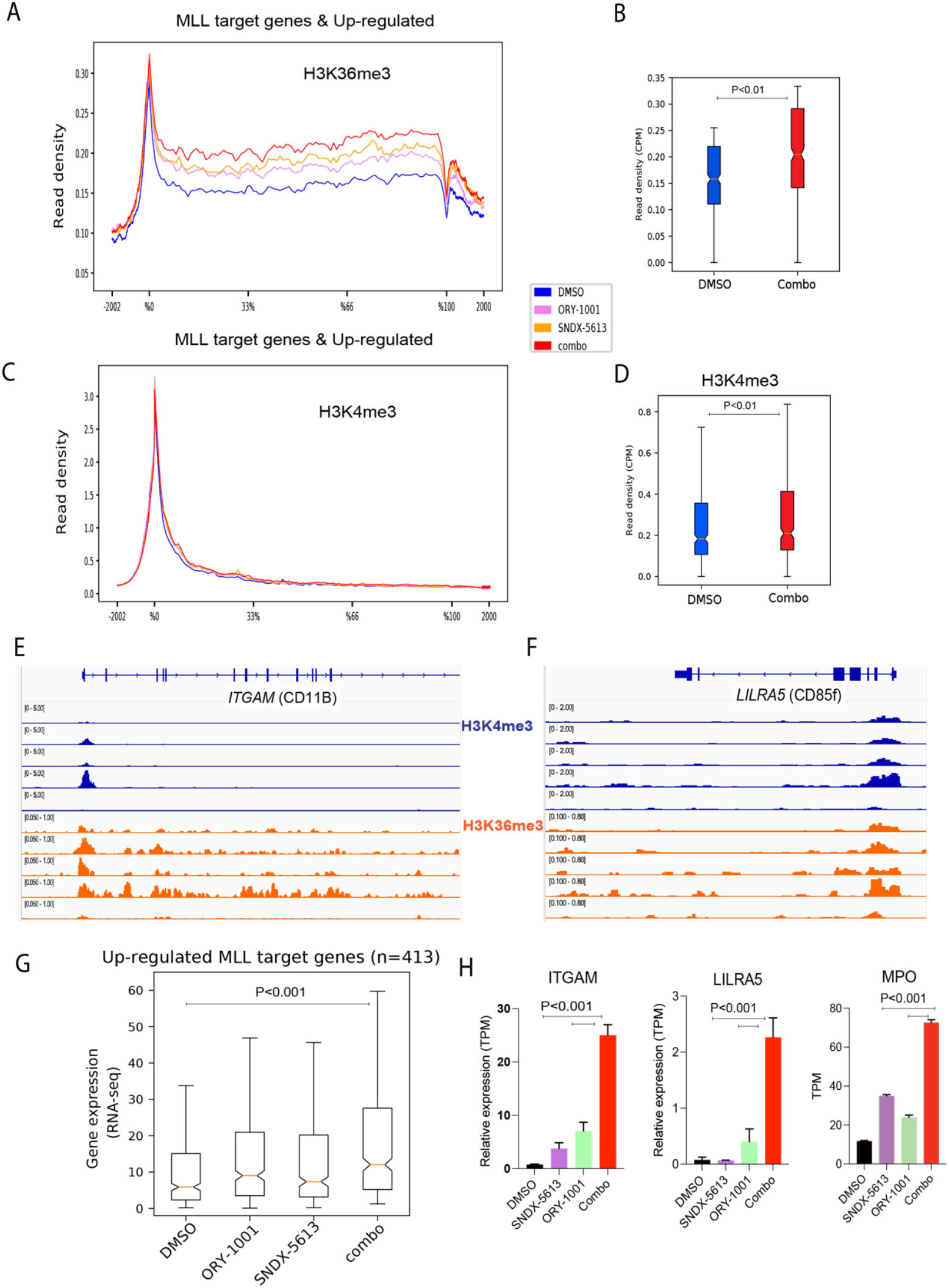
Increases in H3K4me3 at differentiation related genes. **(A-D)** Global metaplot and boxplot of MLL target genes (n = 413), that are upregulated by RNA-seq and associated changes in H3K36me3 and H3K4me3 (**E-F**) Genome tracks at differentiation-related targets (*ITGAM* and *LILRA5* (Leukocyte Immunoglobulin-Like Receptor A5)). **(G)** Box plot of upregulated MLL target genes following treatment with DMSO, SNDX-5613, or the combination treatment. **(H)** TPM (transcripts per million) RNA expression in cells treated with DMSO, SNDX-5613, or the combination treatment (*ITGAM, LILRA5 and MPO* (Myeloperoxidase). Statistical analysis was performed using one-way ANOVA followed by Tukey’s post hoc test (single treatments vs. combination).

The same subset of genes showed an increase in H3K4me3 at their promoters **(Fig. 4C)**, which was further quantified in the box plot demonstrating significant changes after combination treatment **(Fig. 4D)**. These included differentiation-related genes such as *ITGAM* and *LILRA5* (Leukocyte Immunoglobulin Like Receptor A5), as exemplified in **Fig. 4E–F**. The chromatin changes observed in this subset of genes paralleled our RNA-seq data, which showed stepwise upregulation with single-agent treatments and the strongest induction under combination treatment **(Fig. 4G)**. RNA expression of *ITGAM, LILRA5* (examples from Fig. 4D), and *MPO* (*Myeloperoxidase*), an enzyme that generates reactive oxygen species (ROS) and plays a key role in the immune response, showed a marked increase following treatment, consistent with progression toward myeloid differentiation **(Fig. 4H)**.

We next compared these subsets of genes bound by MLL (CUT&RUN) that were either upregulated or downregulated by RNA-seq upon combined LSD1 and menin inhibition **(Fig.S3)**. Interestingly, only the downregulated genes showed substantial progressive loss of MLL occupancy, first modestly with LSD1 inhibition, further with menin inhibition, and most strongly with the combination **(Fig. S3A)**. Interestingly, **Fig. S3** shows that the most prominent changes in the menin-MLL-LEDGF complex and their associated histone marks, H3K4me3 and H3K36me3, occur primarily in the downregulated genes, which are most affected upon combination treatment, consistent with their transcriptional downregulation (Fig. 3F).

For the upregulated MLL-bound subset, menin-MLL-LEDGF occupancy changed only modestly, yet H3K4me3 and H3K36me3 increased markedly (**Fig 4A-F** and **Fig. S3A**). This pattern indicates enhanced functional output of the existing chromatin machinery rather than increased recruitment: higher H3K4me3 is consistent with increased MLL promoter activity, while elevated H3K36me3 marks productive transcriptional elongation (SETD2-dependent) and creates the LEDGF-recognized landscape along gene bodies. Thus, relief of LSD1/CoREST-mediated repression enables the pre-bound menin-MLL-LEDGF axis to operate more effectively, yielding increased H3K4me3/H3K36me3 and transcriptional activation as exemplified by *ITGAM, LILRA5*, and *MPO* in Fig.4H.

### In vivo efficacy of combined Menin and LSD1 inhibition against AML cells

We next evaluated the *in vivo* anti-leukemic efficacy of SNDX-5613 and ORY-1001 in an aggressive and lethal AML patient-derived xenograft (PDX) model expressing MLL-AF6. Following tail-vein infusion and engraftment of cells, mice were treated with vehicle control, SNDX-5613 alone, ORY-1001 alone, or the combination of both inhibitors. The doses used for each drug were previously established as safe [24, 32, 45-47]. Consistent with the *in vitro* findings, co-treatment produced a synergistic therapeutic effect *in vivo*, resulting in a greater reduction of leukemia burden and significantly prolonged survival compared to single-agent treatment in the MLL-AF6 PDX model. The comparison between vehicle- and combination-treated groups demonstrated a marked reduction in AML burden, as indicated by decreased hCD45^+^ cell percentages and improved overall survival, as shown in **Fig. 5**.

**Figure 5.**
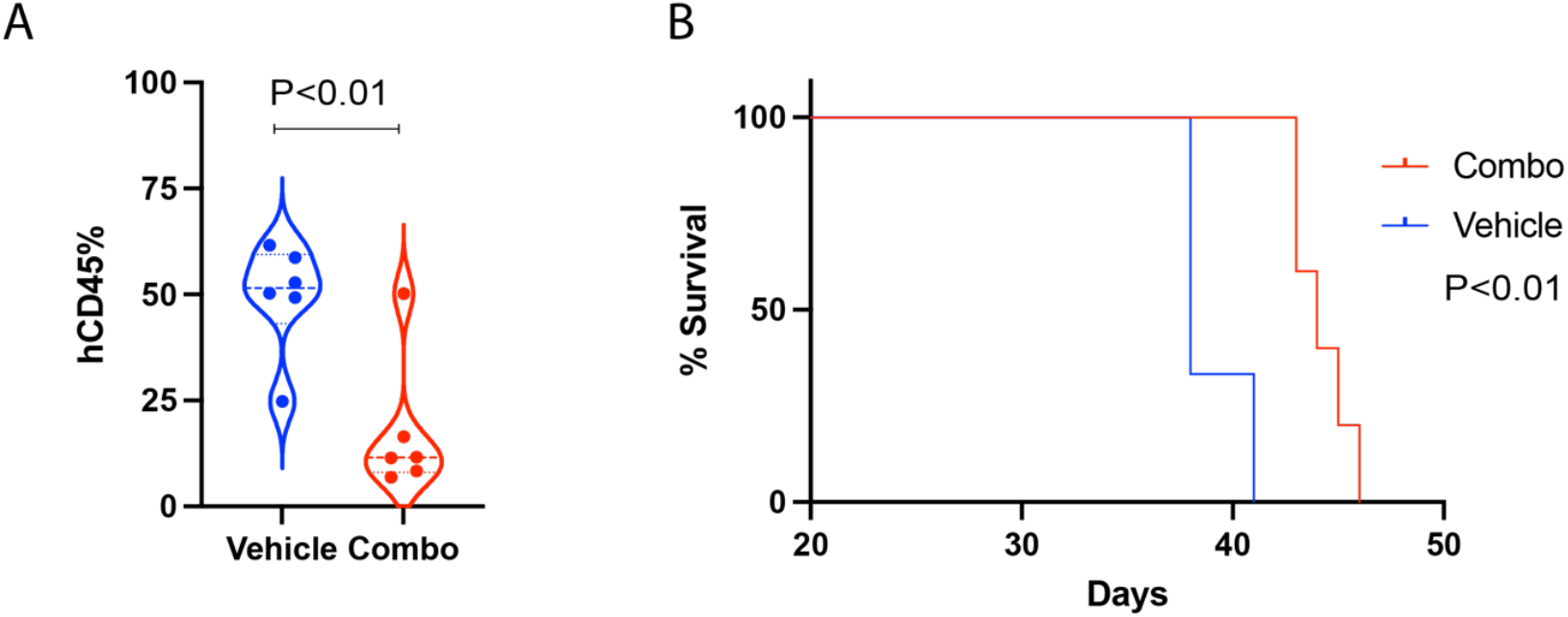
Combination treatment reduces hCD45^+^ leukemic cells and improves survival in MLL-AF6^+^ AML xenografts. **(A)** Flow cytometric analysis of human CD45^+^ (hCD45) cells in bone marrow from vehicle- and combination-treated mice. **(B)** Kaplan-Meier survival curve of NSG mice engrafted with patient-derived MLL-AF6^+^ AML cells in vehicle- and combination-treated mice.

## DISCUSSION

The treatment landscape for MLL-r AML has been transformed by the development of menin inhibitors, which directly disrupt the menin-MLL interaction. Early clinical results have validated this approach, demonstrating meaningful responses in patients with relapsed and refractory disease leading to FDA approval of the first menin inhibitor, revumenib in 2024 [10]. However, as with many targeted therapies, resistance and relapse remain common with monotherapy use [48-50]. This underscores the need for rational drug combinations that not only enhance the depth and durability of response but also address the epigenetic complexity of leukemic transcriptional programs to drive stable differentiation.

By starting with an unbiased epigenetic HTS, we identified LSD1 inhibition as a powerful strategy to potentiate menin blockade. LSD1 has long been appreciated as a transcriptional coregulator and barrier to myeloid differentiation, but its precise mechanistic interplay with the MLL-menin axis had not been defined. Our discovery that LSD1 associates with LEDGF, a PWWP-domain protein central to menin-MLL chromatin tethering, provides new insight into how histone modification readers and erasers cooperate at oncogenic loci. This connection suggests that disrupting both LSD1/CoREST and scaffolding function of menin dismantles a highly integrated chromatin complex essential for leukemic gene expression. The implications extend beyond transcriptional repression of canonical MLL or MLL-r targets. Combination therapy not only diminishes the stability of oncogenic chromatin domains but also shifts the balance toward differentiation. This dual action is particularly relevant given the central role of differentiation blockade in AML pathogenesis. By removing both the sustaining force of MLL-r-driven programs and the epigenetic brake imposed by LSD1, dual inhibition provides a two-pronged mechanism to reprogram leukemic blasts.

From a broader perspective, this work highlights the importance of targeting not only enzymatic activities but also the structural scaffolds that maintain leukemogenic transcriptional hubs. The unexpected binding of LSD1 and LEDGF identified by mass spectrometry following immunoprecipitation, together with their co-occupancy on chromatin and the observed effects on H3K36me3 and H3K4me3 under combination treatment, suggests that disrupting these higher-order interactions is more critical than inhibiting the individual factors themselves. Such insights may extend to other fusion-driven malignancies where reader-eraser-scaffold partnerships reinforce transcriptional addiction. The mechanistic basis of increased LSD1 occupancy upon menin inhibition also warrants further investigation, as it may represent either a compensatory response or a novel vulnerability. In the combination treatment, LSD1 is already inhibited, which may lead to its dissociation from chromatin as part of the corepressor complex. As a result, LSD1 can no longer effectively repress these loci. This could explain why, in certain downregulated gene subsets, the effect of the combination is less pronounced compared to the strong activation observed in the upregulated genes.

Preclinical in vivo studies demonstrated that dual inhibition is effective and tolerable, even in patient-derived xenografts, suggesting a therapeutic window compatible with clinical development. The observation that toxicity was limited and transient, while survival was significantly improved, reinforces the feasibility of this approach. These findings are timely, given emerging reports of menin inhibitor resistance mediated by secondary mutations, adaptive chromatin reorganization, or lineage plasticity. LSD1 co-inhibition may mitigate such mechanisms by disrupting chromatin anchoring and restoring differentiation pathways, thereby reducing opportunities for therapeutic escape. Future studies should also assess whether combined inhibition can prevent or overcome acquired resistance to menin inhibitors in patients. Finally, clinical translation will require careful exploration of dosing, scheduling, and potential interactions with existing therapies, as well as the identification of biomarkers to guide patient selection.

Collectively, our findings establish LSD1 as a critical cofactor within the menin-MLL transcriptional complex through its interaction with LEDGF and reveal that combined inhibition of menin and LSD1 exerts complementary and synergistic effects. This dual blockade dismantles MLL-r-driven transcriptional programs, changes critical histone modifications, promotes differentiation, and translates into potent in vivo efficacy. Given the limitations of menin inhibitor monotherapy, our work provides a strong mechanistic rationale for clinical evaluation of menin and LSD1 inhibitor combinations in MLL-r AML.

## METHODS

### Cell viability assays

Cells were seeded at a density of 2 × 10^4^ cells/mL in 96-well plates (Greiner Bio-One, Cat# 655180). Treatments included DMSO (vehicle control) and serial dilutions of the test compounds for 4 days. Each treatment condition was performed with five technical replicates per concentration, and the entire experiment was repeated in three independent biological replicates.

### Apoptosis analysis

The AML cells were seeded (2 × 10^5^ cells/mL) and treated with ORY-1001 and SNDX-5613 (purchased from MedChemExpress (MCE)) at the final concentrations of 0.2 μM for 4 days. Then, the AML cells were harvested and stained with Annexin V-PE and 7-AAD according to BD’s instructions (BD, USA), followed by apoptosis analysis by flow cytometry.

### Flow cytometry

AML cell lines were treated with ORY-1001 and/or SNDX-5613 (MCE) for 4 days. For surface immunophenotyping, 5 × 10^5 cells were stained for CD11b using PE-conjugated anti-human CD11b (BD Pharmingen, #555388), following the manufacturer’s instructions. Data were analyzed after gating on singlet, viable cells; fluorescence compensation and instrument settings were kept constant across conditions.

### Mass Spectrometry Analysis

Following immunoprecipitation of endogenous LSD1 in MOLM-13, final eluates were subjected to mass spectrometry analysis (n = 2 biological replicates per condition). Proteins were digested in-solution with trypsin, and peptides were purified using C18 spin columns. Peptide mixtures were analyzed using a 1.5-hour liquid chromatography gradient on a Thermo Q Exactive HF or Orbitrap Exactive HF mass spectrometer. Raw MS data were processed in MaxQuant (version 1.6.17.0) against the UniProt Human database (release 07/21/2022 [51]) using MaxQuant 1.6.17.0. [52]. Searches were performed with full tryptic specificity and included variable modifications for N-terminal acetylation, methionine oxidation, and asparagine deamidation. Protein and peptide false discovery rates (FDR) were set to 1%. Label-free quantification (LFQ) values were normalized using the MaxLFQ algorithm [53]. Proteomic data analysis was conducted in Perseus (Tyanova, Temu, et al., 2016) following standard procedures for label-free interaction profiling. Reverse hits, site-only identifications, and common contaminants were excluded. Intensity-Based Absolute Quantification (iBAQ) intensities were log2-transformed, and proteins were retained if at least two non-zero values were present in either the IP or control groups (n = 2 per group). Missing values were imputed using a normal distribution (width = 0.3, downshift = 1.8). Statistically significant interactors were identified using a two-sample t-test with an s_0_ parameter of 2 and a permutation-based FDR threshold of <0.001.

### CUT&RUN Assay and Sequencing

CUT&RUN was performed using the CUTANA™ ChIC/CUT&RUN Kit (EpiCypher, Cat. No. 14-1048) according to the manufacturer’s protocol. Briefly, MOLM-13 cells were treated for 24 hours, after which 5 × 10^5^ cells were harvested, permeabilized with digitonin, and assessed using Trypan blue staining. Cells were bound to activated Concanavalin A beads and incubated overnight at 4 °C with primary antibodies against H3K4me3 (EpiCypher, SKU: 13-0060), H3K36me3 (EpiCypher, SKU: 13-0058), H3K27me3 (EpiCypher, SKU: 13-0055), LSD1

(Abcam, ab129195), menin (EpiCypher, SKU: 13-2021), MLL1 (KMT2A) (EpiCypher, SKU: 13-2004), LEDGF (Catalog # A300-848A) or IgG control (EpiCypher, SKU: 13-0042K). Following washes, cells were incubated with pAG-MNase (EpiCypher, SKU: 15-1016) for 10 minutes at room temperature, and chromatin was digested for 2 hours at 4 °C. Reactions were stopped with stop buffer supplemented with E. coli spike-in DNA (EpiCypher, SKU: 18-1401), and DNA fragments were purified with DNA cleanup columns. Sequencing libraries were prepared using the CUTANA™ DNA Library Prep Kit (EpiCypher, SKU: 14-1001) and sequenced on an Illumina NovaSeq 6000, generating at least 10 million paired-end reads per sample (150 bp).

### CUT&RUN Analysis

Sequencing reads were processed using the nf-core/cutandrun pipeline (v3.1, doi: 10.5281/zenodo.5653535) [54, 55], on the Pluto bioinformatics platform (https://pluto.bio). Adapter sequences were trimmed with Trim Galore, and reads were aligned to the human GRCh38 reference genome (NCBI build p.14, release 110) using Bowtie2. Duplicate reads were removed with Picard. Peak calling was performed with MACS2 using CPM normalization, defining peaks as regions where signal in target samples exceeded IgG controls. Consensus peaks across all samples were merged with BEDTools and annotated to the nearest transcription start site (TSS) using HOMER. Peak counts were quantified with featureCounts for downstream analyses.

### RNA Sequencing (RNA-seq)

For RNA-seq, 500 ng of total RNA was extracted using TRIzol (Thermo Fisher Scientific, Cat. #15596026), treated with DNase I (RNase-free, Cat. #AM1907), and processed with the TruSeq Stranded Total RNA Library Prep Kit (Illumina, Cat. #20020596). Libraries were sequenced on an Illumina NovaSeq 6000, generating a minimum of 50 million paired-end reads per sample (100 bp).

### RNA-seq Analysis

FASTQ files were trimmed for adapters and low-quality bases using Trimmomatic (RRID:SCR_011848) [56]. Reads were aligned to the hg38 human genome with STAR aligner (RRID:SCR_015899) using default parameters. Gene-level quantification was performed with RSEM (RRID:SCR_013027) [57] against the Ensembl reference transcriptome (release 97) settings [58]. with expression normalized to transcripts per million (TPM). Differential expression analysis was performed with DESeq2, applying a 1.5-fold change cutoff and FDR < 0.05 unless otherwise specified. Heatmaps were generated with clustermap (Ward’s method; Euclidean distance; https://doi.org/10.21105/joss.03021) to visualize Z-score row-normalized gene expression and clustering patterns. Functional enrichment of differentially expressed genes was assessed using Enrichr [59] against GO Biological Process terms.

### ATAC-seq and Data Analysis

ATAC-seq was performed following established protocols [60] with minor modifications. Briefly, DNA libraries were quantified using the Qubit dsDNA High Sensitivity kit (Thermo Fisher Scientific) on a Qubit 2 fluorometer, pooled, and sequenced on an Illumina NovaSeq 6000 to generate paired-end 100 bp reads. Raw sequencing data were processed as described in [61].

### In vivo studies

All animal studies were performed under approval of the University of Miami Institutional Animal Care and Use Committee (IACUC). We obtained the MLL-AF6 PDX model (CPDM-0984X) from the Dana Farber Cancer Institute. We passaged the tumor in vivo in SGM3-NOD scid gamma (SGM3-NSG) mice (Jackson Laboratories #: 013062) through tail-vein injection of 2 million cells per animal. For the efficacy experiment, 12 week female mice were pre-conditioned with 1Gy the same day of the tail-vein tumor injection. We determined tumor burden by flow cytometric analysis of human CD45 in peripheral blood, with body weight loss >20% as a predetermined survival endpoint. SNDX-5613 (purchased from MedChemExpress (MCE)) treatment was at 40 mg/kg twice daily (5 days on, 2 days off) by oral gavage. ORY-1001 (purchased from MCE) treatment was at 0.02 mg/kg once daily three times a week by oral gavage. Both drugs were prepared in a mix of 10% DMSO, 40% PEG300, 5% Tween-80, and 45% saline.

## Supporting information

Figure S1. ATAC-seq analysis demonstrating synergistic increased chromatin accessibility with combination treatment compared to either single agent in

Figure S2. (A-C) Genome browser tracks at the canonical MLL target genes Meis1, HOXA cluster and MYC loci showing occupancy of histone modifications (

Figure S3. MLL target genes that are upregulated (dark blue) or downregulated (green) display distinct binding pattern to MLL-menin-LEDGF complex (A)

## Data Availability Statement

All data supporting the findings of this study are available within the main text and/or the Supplementary Materials. Raw and processed RNA-seq, ATAC-seq and CUT&RUN datasets, as well as relevant data processing scripts have been deposited on University of Miami Dataverse.

## AUTHOR CONTRIBUTIONS

**M.M. Tayari** Led the conceptualization and execution of the study. Performed experiments including IC_50_ and cell viability assays and synergy assays, RNA-seq, ATAC-seq, CUT&RUN, western blotting, and endogenous co-immunoprecipitations. Conducted formal data analysis using Pluto and GraphPad Prism, and contributed to validation, investigation, visualization, methodology, original draft writing, as well as review and editing of the manuscript. **H. Gomes Dos Santos, F. Beckedorf, G.M. Lavezzo** and **M.M. Tayari** performed bioinformatics analyses. **T. Totiger, A. Mookhtiar** and **A.L. Kingham** prepared samples for epigenetic screening, performed cytospin and apoptosis assay. **G. Mas Martin, H-T, Huang, E. Karaca, D. Bilbao** performed in vivo testing and analysis. **J.Taylor, J.M. Watts** and **R. Shiekhattar**: oversaw project conceptualization, supported the project through resources, validation, investigation, and project administration, and participated in reviewing and editing the manuscript.

## ACKNOWLEDGEMENTS

We are grateful to the members of the Shiekhattar lab for their valuable discussions and feedback. We thank the Proteomics and Metabolomics Facility at the Wistar Institute for mass spectrometry analysis. This work was supported by funding from the University of Miami Miller School of Medicine, the Sylvester Comprehensive Cancer Center, and NIH grant R01GM078455 awarded to R.S. Research reported in this publication was also supported by the Cancer Modeling Shared Resource (CMSR; RRID: SCR_022891), the Oncogenomics Shared Resource (OGSR; SCR_022502) of the Sylvester Comprehensive Cancer Center at the University of Miami, which are supported by the National Cancer Institute (NCI) of the National Institutes of Health (NIH) under award number P30CA240139. The content is solely the responsibility of the authors and does not necessarily represent the official views of the NIH. This work was further supported by the Leukemia & Lymphoma Society’s Specialized Center of Research Program (LLS-SCOR), awarded to Stephen D. Nimer (Program PI), with support for Project 1 to R. Shiekhattar and J.M. Watts.

## REFERENCES

1. Bernt, K.M. and S.A. Armstrong, Targeting epigenetic programs in MLL-rearranged leukemias. Hematology Am Soc Hematol Educ Program, 2011. 2011: p. 354–60.

2. Krivtsov, A.V. and S.A. Armstrong, MLL translocations, histone modifications and leukaemia stem-cell development. Nat Rev Cancer, 2007. 7(11): p. 823–33.

3. Slany, R.K., the molecular biology of mixed lineage leukemia. Haematologica, 2009. 94(7): p. 984–93.

4. Meyer, C., et al., The MLL recombinome of acute leukemias in 2017. Leukemia, 2018. 32(2): p. 273–284.

5. Meyer, C., et al., New insights to the MLL recombinome of acute leukemias. Leukemia, 2009. 23(8) p. 1490–9.

6. Charles, N.J. and D.F. Boyer, Mixed-Phenotype Acute Leukemia: Diagnostic Criteria and PiKalls. Arch Pathol Lab Med, 2017. 141(11): p. 1462–1468.

7. Krivtsov, A.V., et al., A Menin-MLL Inhibitor Induces Specific Chromatin Changes and Eradicates Disease in Models of MLL-Rearranged Leukemia. Cancer Cell, 2019. 36(6): p. 660–673.e11.

8. Marschalek, R., MLL leukemia and future treatment strategies. Arch Pharm (Weinheim), 2015. 348(4): p. 221–8.

9. Winters, A.C. and K.M. Bernt, MLL-Rearranged Leukemias-An Update on Science and Clinical Approaches. Front Pediatr, 2017. 5: p. 4.

10. Aldoss, I., et al., Updated Results and Longer Follow-up from the AUGMENT-101 Phase 2 Study of Revumenib in All Patients with Relapsed or Refractory (R/R) KMT2Ar Acute Leukemia. Blood, 2024. 144(Supplement 1): p. 211–211.

11. Yokoyama, A. and M.L. Cleary, Menin critically links MLL proteins with LEDGF on cancer-associated target genes. Cancer Cell, 2008. 14(1): p. 36–46.

12. Zhang, Y., et al., Disordered epigenetic regulation in MLL-related leukemia. Int J Hematol, 2012. 96(4): p. 428–37.

13. Huang, J., et al., The same pocket in menin binds both MLL and JUND but has opposite effects on transcription. Nature, 2012. 482(7386): p. 542–6.

14. El Ashkar, S., et al., LEDGF/p75 is dispensable for hematopoiesis but essential for MLL-rearranged leukemogenesis. Blood, 2018. 131(1): p. 95–107.

15. Sharma, S., et al., Affinity switching of the LEDGF/p75 IBD interactome is governed by kinase-dependent phosphorylation. Proc Natl Acad Sci U S A, 2018. 115(30): p. E7053–e7062.

16. Eidahl, J.O., et al., Structural basis for high-affinity binding of LEDGF PWWP to mononucleosomes. Nucleic Acids Res, 2013. 41(6): p. 3924–36.

17. Pradeepa, M.M., et al., Psip1/Ledgf p75 restrains Hox gene expression by recruiting both trithorax and polycomb group proteins. Nucleic Acids Res, 2014. 42(14): p. 9021–32.

18. LeRoy, G., et al., LEDGF and HDGF2 relieve the nucleosome-induced barrier to transcription in differentiated cells. Sci Adv, 2019. 5(10): p. eaay3068.

19. Čermáková, K., et al., Validation and Structural Characterization of the LEDGF/p75–MLL Interface as a New Target for the Treatment of MLL-Dependent Leukemia. Cancer Research, 2014. 74(18): p. 5139–5151.

20. Kühn, M.W., et al., Targeting Chromatin Regulators Inhibits Leukemogenic Gene Expression in NPM1 Mutant Leukemia. Cancer Discov, 2016. 6(10): p. 1166–1181.

21. Klossowski, S., et al., Menin inhibitor MI-3454 induces remission in MLL1-rearranged and NPM1-mutated models of leukemia. J Clin Invest, 2020. 130(2): p. 981–997.

22. Issa, G.C., et al., The menin inhibitor revumenib in KMT2A-rearranged or NPM1-mutant leukaemia. Nature, 2023. 615(7954): p. 920–924.

23. Issa, G.C., et al., Menin Inhibition With Revumenib for KMT2A-Rearranged Relapsed or Refractory Acute Leukemia (AUGMENT-101). J Clin Oncol, 2025. 43(1): p. 75–84.

24. Arellano, M.L., et al., Menin inhibition with revumenib for NPM1-mutated relapsed or refractory acute myeloid leukemia: the AUGMENT-101 study. Blood, 2025. 146(9): p. 1065–1077.

25. Lee, M.G., et al., An essential role for CoREST in nucleosomal histone 3 lysine 4 demethylation. Nature, 2005. 437(7057): p. 432–5.

26. Shi, Y., et al., Histone demethylation mediated by the nuclear amine oxidase homolog LSD1. Cell, 2004. 119(7): p.941–53.

27. Harris, W.J., et al., The histone demethylase KDM1A sustains the oncogenic potential of MLL-AF9 leukemia stem cells. Cancer Cell, 2012. 21(4): p. 473–87.

28. Maiques-Diaz, A., et al., Enhancer Activation by Pharmacologic Displacement of LSD1 from GFI1 Induces Differentiation in Acute Myeloid Leukemia. Cell Rep, 2018. 22(13): p. 3641–3659.

29. Ravasio, R., et al., Targeting the scaffolding role of LSD1 (KDM1A) poises acute myeloid leukemia cells for retinoic acid-induced differentiation. Sci Adv, 2020. 6(15): p. eaax2746.

30. Tayari, M.M., et al., Clinical Responsiveness to All-trans Retinoic Acid Is Potentiated by LSD1 Inhibition and Associated with a Quiescent Transcriptome in Myeloid Malignancies. Clin Cancer Res, 2021. 27(7): p.1893–1903.

31. Wass, M., et al., A proof of concept phase I/II pilot trial of LSD1 inhibition by tranylcypromine combined with ATRA in refractory/relapsed AML patients not eligible for intensive therapy. Leukemia, 2021. 35(3): p. 701–711.

32. Salamero, O., et al., First-in-Human Phase I Study of Iadademstat (ORY-1001): A First-in-Class Lysine-Specific Histone Demethylase 1A Inhibitor, in Relapsed or Refractory Acute Myeloid Leukemia. J Clin Oncol, 2020. 38(36): p. 4260–4273.

33. Adriaanse, F.R.S., et al., Distinct Responses to Menin Inhibition and Synergy with DOT1L Inhibition in KMT2A-Rearranged Acute Lymphoblastic and Myeloid Leukemia. Internaeonal Journal of Molecular Sciences, 2024. 25(11): p. 6020.

34. Olsen, S.N., et al., Combined inhibition of KAT6A/B and Menin reverses estrogen receptor-driven gene expression programs in breast cancer. Cell Rep Med, 2025. 6(7): p. 102192.

35. Issa, G.C., et al., Combination Strategies with Menin Inhibitors for Acute Leukemia. Blood Cancer Discov, 2025.

36. Hakimi, M.A., et al., A core-BRAF35 complex containing histone deacetylase mediates repression of neuronal-specific genes. Proc Natl Acad Sci U S A, 2002. 99(11): p. 7420–5.

37. Hakimi, M.A., et al., A candidate X-linked mental retardation gene is a component of a new family of histone deacetylase-containing complexes. J Biol Chem, 2003. 278(9): p. 7234–9.

38. Barrios Á, P., et al., Differential properties of transcriptional complexes formed by the CoREST family. Mol Cell Biol, 2014. 34(14): p. 2760–70.

39. Saleque, S., et al., Epigenetic regulation of hematopoietic differentiation by Gfi-1 and Gfi-1b is mediated by the cofactors CoREST and LSD1. Mol Cell, 2007. 27(4): p. 562–72.

40. Nicosia, L., et al., Pharmacological inhibition of LSD1 triggers myeloid differentiation by targeting GSE1 oncogenic functions in AML. Oncogene, 2022. 41(6): p. 878–894.

41. Vcelkova, T., et al., GSE1 links the HDAC1/CoREST co-repressor complex to DNA damage. Nucleic Acids Res, 2023. 51(21): p. 11748–11769.

42. Liu, J., et al., Arginine methylation-dependent LSD1 stability promotes invasion and metastasis of breast cancer. EMBO Rep, 2020. 21(2): p. e48597.

43. Schenk, T., et al., Inhibition of the LSD1 (KDM1A) demethylase reactivates the all-trans-retinoic acid differentiation pathway in acute myeloid leukemia. Nat Med, 2012. 18(4): p. 605–11.

44. Hosseini, A., et al., Perturbing LSD1 and WNT rewires transcription to synergistically induce AML differentiation. Nature, 2025. 642(8067): p. 508–518.

45. Fiskus, W., et al., Targeting of epigenetic co-dependencies enhances anti-AML efficacy of Menin inhibitor in AML with MLL1-r or mutant NPM1. Blood Cancer J, 2023. 13(1): p. 53.

46. Lynch, E.J., et al., Revumenib for Relapsed or Refractory Acute Leukemia With a KMT2A Translocation. Ann Pharmacother, 2025: p. 10600280251341279.

47. Salamero, O., et al., Iadademstat in combination with azacitidine in patients with newly diagnosed acute myeloid leukaemia (ALICE): an open-label, phase 2a dose-finding study. Lancet Haematol, 2024. 11(7): p. e487–e498.

48. Perner, F., et al., MEN1 mutations mediate clinical resistance to menin inhibition. Nature, 2023. 615(7954): p. 913–919.

49. Boussi, L., S.F. Cai, and E.M. Stein, Advances in menin inhibition in acute myeloid leukemia. Trends in Cancer, 2025. 11(9): p. 889–900.

50. Zhou, X., et al., Decoding the Epigenetic Drivers of Menin-MLL Inhibitor Resistance in KMT2A-Rearranged Acute Myeloid Leukemia. Blood, 2023. 142: p. 587.

51. Bowler-Barnei, E.H., et al., UniProt and Mass Spectrometry-Based Proteomics-A 2-Way Working Relationship. Mol Cell Proteomics, 2023. 22(8): p. 100591.

52. Tyanova, S., T. Temu, and J. Cox, The MaxQuant computational plaKorm for mass spectrometry-based shotgun proteomics. Nature Protocols, 2016. 11(12): p. 2301–2319.

53. Cox, J., et al., Accurate proteome-wide label-free quantification by delayed normalization and maximal peptide ratio extraction, termed MaxLFQ. Mol Cell Proteomics, 2014. 13(9): p. 2513–26.

54. Ewels, P.A., et al., The nf-core framework for community-curated bioinformatics pipelines. Nat Biotechnol, 2020. 38(3): p. 276–278.

55. Di Tommaso, P., et al., NexKlow enables reproducible computational workfi ows. Nature Biotechnology, 2017. 35(4): p. 316–319.

56. Bolger, A.M., M. Lohse, and B. Usadel, Trimmomatic: a fi exible trimmer for Illumina sequence data. Bioinformaecs, 2014. 30(15): p. 2114–20.

57. Langmead, B., et al., Ultrafast and memory-efficient alignment of short DNA sequences to the human genome. Genome Biol, 2009. 10(3): p. R25.

58. Li, B. and C.N. Dewey, RSEM: accurate transcript quantification from RNA-Seq data with or without a reference genome. BMC Bioinformaecs, 2011. 12: p. 323.

59. Chen, E.Y., et al., Enrichr: interactive and collaborative HTML5 gene list enrichment analysis tool. BMC Bioinformaecs, 2013. 14: p. 128.

60. Corces, M.R., et al., An improved ATAC-seq protocol reduces background and enables interrogation of frozen tissues. Nat Methods, 2017. 14(10): p. 959–962.

61. Chan, H.L., et al., Polycomb complexes associate with enhancers and promote oncogenic transcriptional programs in cancer through multiple mechanisms. Nat Commun, 2018. 9(1): p. 3377.

