## Supplementary figures and images for "Co-targeting menin and LSD1 dismantles oncogenic programs and restores differentiation in MLL-rearranged AML"

### Figure S1. ATAC-seq analysis demonstrating synergistic increased chromatin accessibility with combination treatment compared to either single agent in

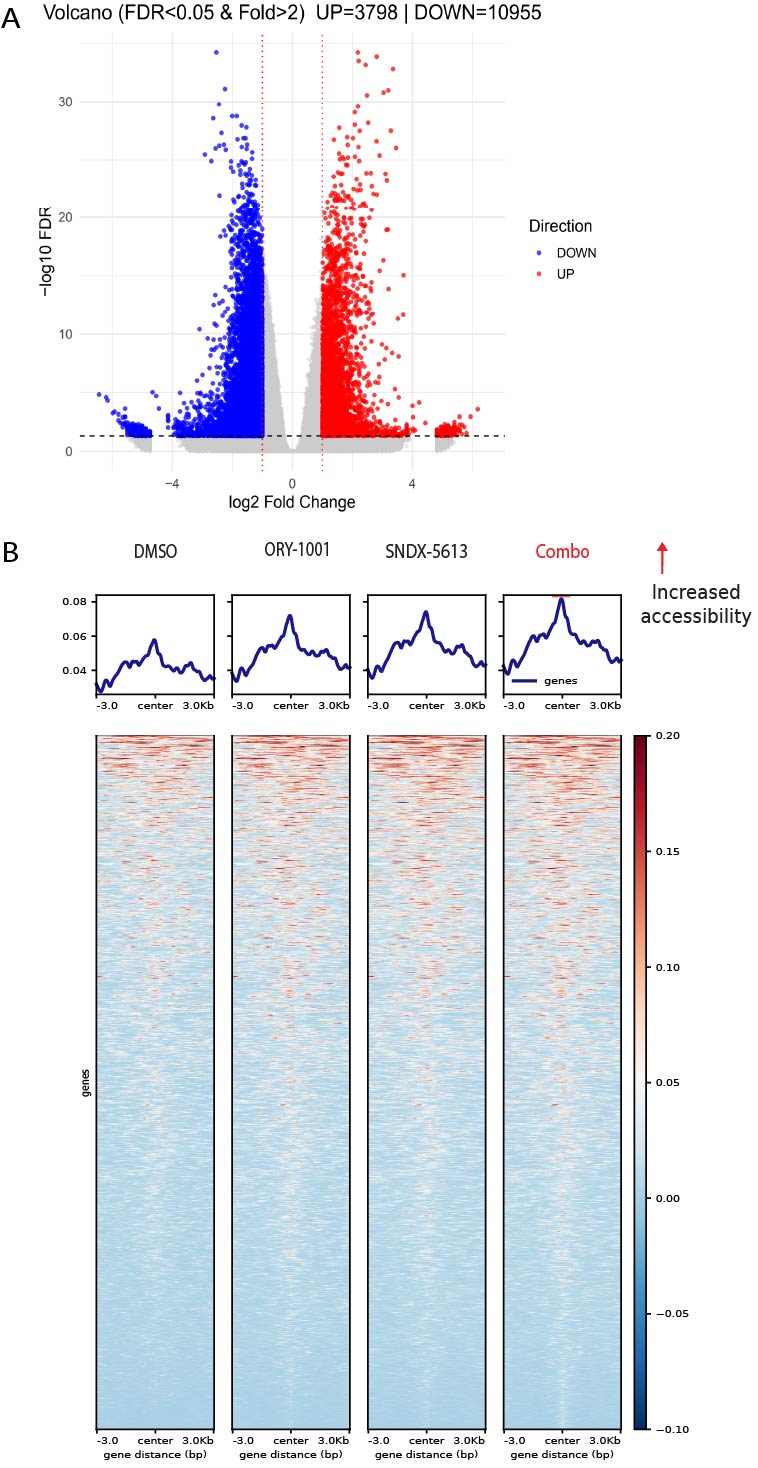

### Figure S2. (A-C) Genome browser tracks at the canonical MLL target genes Meis1, HOXA cluster and MYC loci showing occupancy of histone modifications (

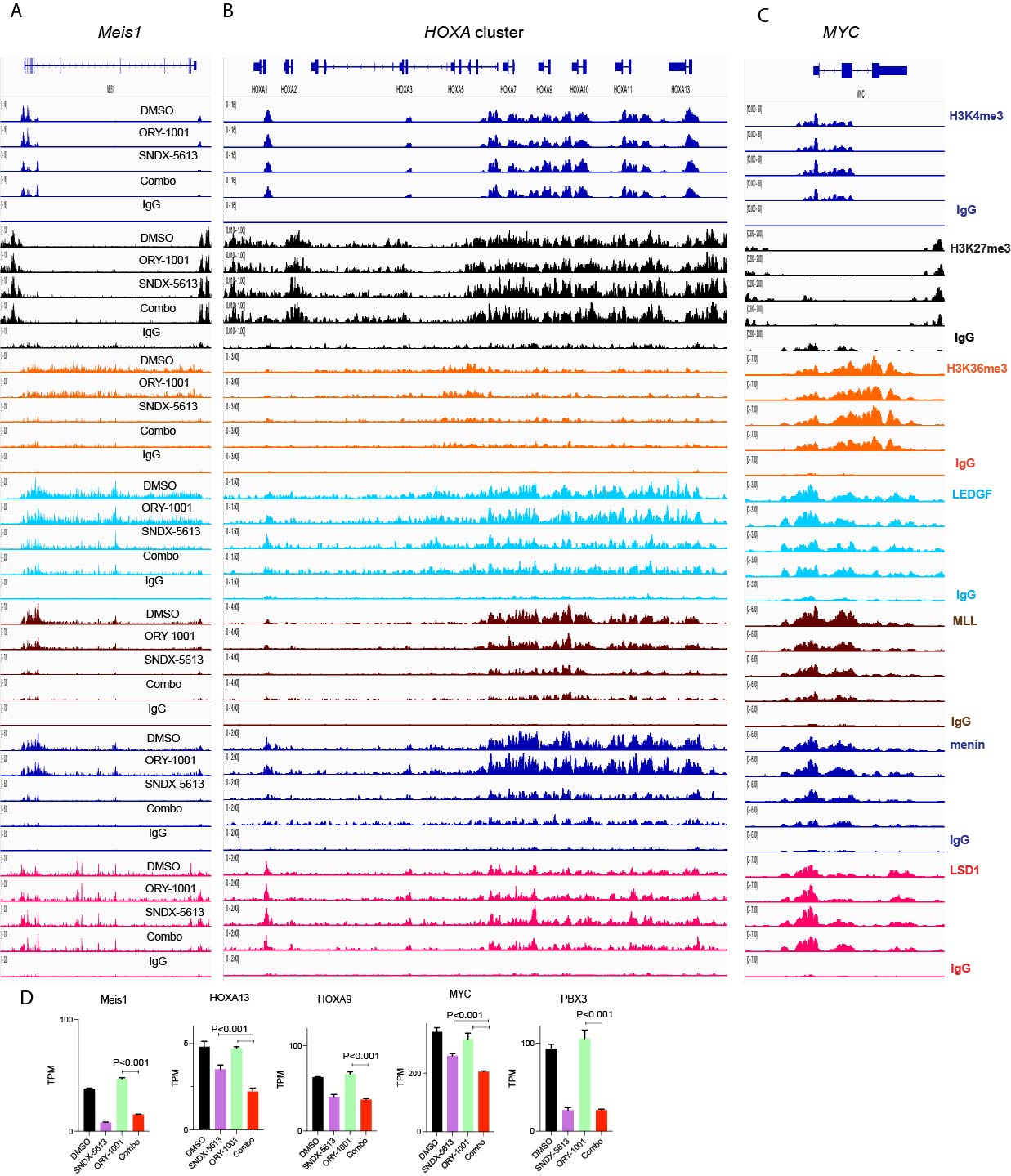

### Figure S3. MLL target genes that are upregulated (dark blue) or downregulated (green) display distinct binding pattern to MLL-menin-LEDGF complex (A)

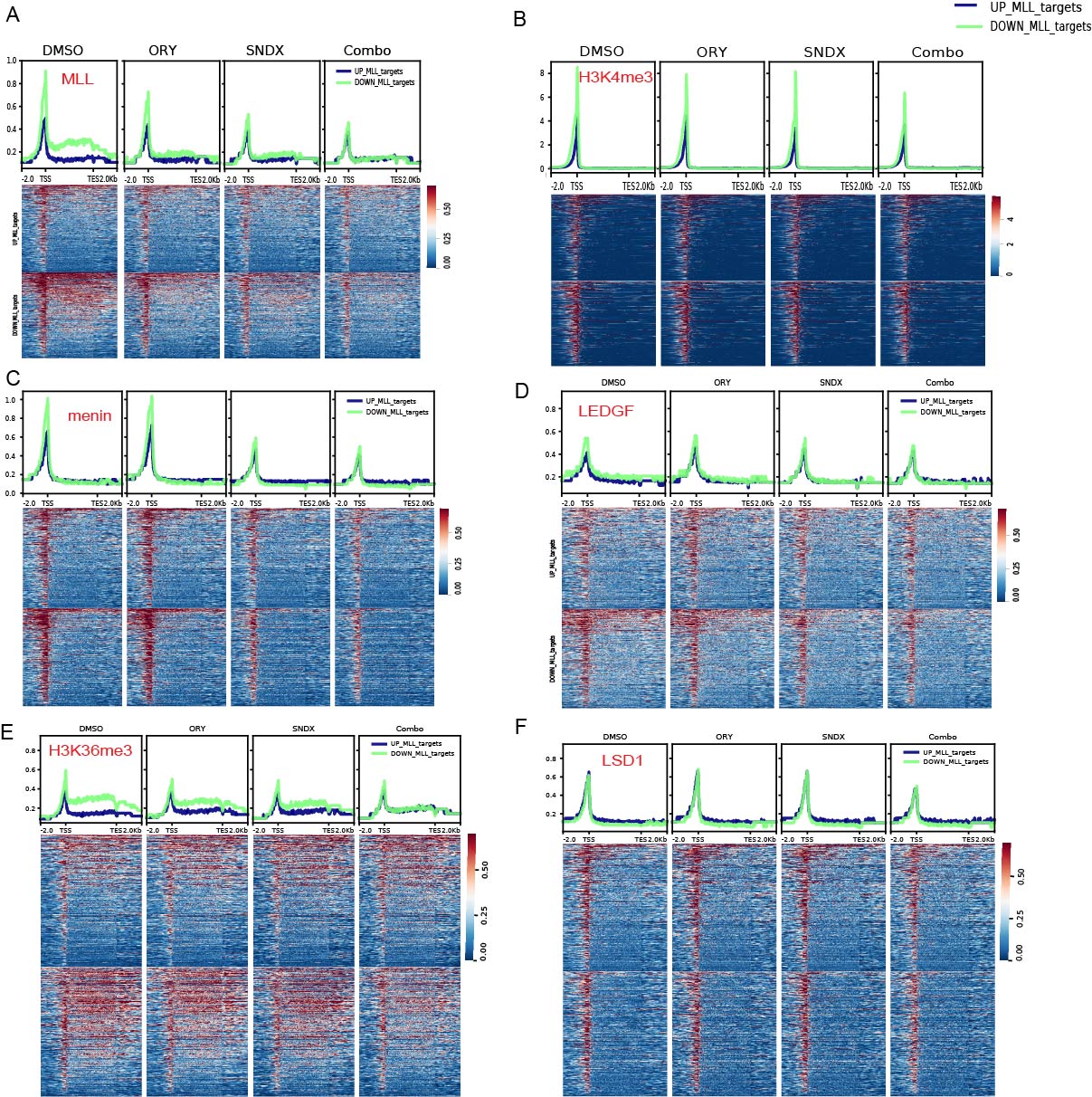
